# Customizable host and viral transcript enrichment using CRISPR-Cas9 long-read sequencing for isoform discovery and validation

**DOI:** 10.1101/2025.04.11.648353

**Authors:** An N.T. Nguyen, Jianshu Zhang, Shuxin Zhang, Miranda E. Pitt, Devika Ganesamoorthy, Svenja Fritzlar, Jessie J.-Y. Chang, Sarah L. Londrigan, Lachlan J.M. Coin

## Abstract

Long-read RNA sequencing has been broadly utilized to examine the diversity of transcriptomes, understand differential expression and discover novel transcript isoforms. One of the major limitations of whole transcriptome sequencing is the difficulty in obtaining sufficient depth for low abundant transcripts. Methods which address this are either difficult to scale or customize: long-range PCR is customizable but difficult to scale beyond a few targets; probe hybridization panels are suited for scaling but require substantial investment to customize. In this study, we adopted RNA-guided CRISPR-Cas9 nuclease-based enrichment to target specific human and SARS-CoV-2 transcripts followed by long-read sequencing, utilizing minimal number of guide RNAs per target isoform. Our findings demonstrate that the CRISPR-Cas system is a highly effective method for customizable long-read sequencing of target transcripts while maintaining the accuracy of relative gene expression levels. The results highlight a valuable method for future research on transcript enrichment for isoform identification and low abundance transcript detection in infectious disease diagnosis.

## Main text

In eukaryotic cells, a single gene can produce multiple messenger RNA (mRNA) products through cell type-specific processes such as alternative transcription initiation, splicing, intron retention and polyadenylation^1-3^. In humans, approximately 95% of all protein-coding genes undergo alternative splicing to varying degrees^4^. Each transcript isoform can contain a unique set of features that influence protein production differently^5^, thereby diversifying the transcriptome and proteome. Switching between transcript isoforms have been detected in several diseases and infections^6-8^, including infections with Severe Acute Respiratory Coronavirus 2 (SARS-CoV-2)^9^. Thus, examining the transcriptome diversity not only at the gene level but also at the isoform level is necessary for fully understanding disease pathogenesis.

Long-read RNA sequencing has been broadly used to understand gene expression in different cell types and to resolve full-length transcript isoforms as well as other complex RNA processing events^10-13^. Pacific Biosciences (PacBio)^14^ and Oxford Nanopore Technologies (ONT)^15^ are two dominant platforms that have overcome the major limitations caused by short-read sequencing by generating reads longer than 10 kb and directly sequencing full-length transcript molecules end-to-end instead of short reads at 150 - 500 bp^16,17^. ONT sequencing platform provides two PCR-free protocols to generate long-read RNA sequencing, direct cDNA sequencing and direct RNA sequencing (dRNA-seq). While direct cDNA sequencing requires reverse transcription, dRNA-seq allows native RNA molecules to be translocated through nanopores, avoiding both reverse transcription and PCR amplification^18^. However, the process often involves the sequencing of the whole transcriptome, leading to incomplete assessment of isoform diversity as well as inability to accurately estimate isoform proportions, particularly for low-abundance transcripts. Moreover, many diagnostic assays only require information on expression of a subset of transcripts. Several targeted sequencing approaches have been developed and allow deeper sequencing of specific transcripts within the complex transcriptomes. Pre-amplifying the target transcripts by using long-range RT-PCR followed by long-read sequencing can generate full-length transcripts. Yet, primer cross-reactivity and amplification bias lead to the difficulty of scaling up to large gene panels^19^. Hybridization capture-based enrichment using biotinylated^20,21^ and synthesizing capture oligos^22^ have been recently introduced. These methods rely on generation of oligos with high tiling density across target transcripts, which is either expensive when ordered from commercial supplier, or requires complex experimental protocols to create. Moreover, adding or subtracting transcripts from an existing panel is not straightforward. Finally, the hybridization workflows can be time consuming.

Clustered Regularly Interspaced Short Palindromic Repeats (CRISPR)/ CRISPR-associated protein 9 (Cas9) is a powerful genome editing and cloning tool^23^. ONT long-read sequencing platform first adapted CRISPR-Cas9 to enrich chromosomal DNA and introduced CATCH (Cas9-assisted targeting of chromosome segments) sequencing^24^. Cas9 nuclease cleaves the regions of interest which are then separated by size selection using by pulsed field gel electrophoresis (PFGE). However, this approach generates biases due to the follow-up PCR amplification of the target to compensate for the low recovery. Later, nanopore Cas9-targeted sequencing (nCAT) directly ligates adapters to Cas9-cleaved DNA products^25^. This method has successfully enriched several DNA targets. However, the potential of applying this method to enrich RNA transcripts has not yet been explored.

Here, we employed Cas9-guided targeted sequencing to enrich specific RNA transcripts within complex transcriptomes of different human cell lines with and without SARS-CoV-2 infections. We first validated the method on five human genes with a wide range of expression levels. We then extended the enrichment strategy to SARS-CoV-2 infected cells with low viral loads to detect viral transcripts and identify the expression transcript isoforms of human cells with the infection.

## Results

### High enrichment of target transcripts by CRISPR-Cas9 targeted sequencing

To investigate the enrichment efficiency, we first chose five human genes encoded for *FTL* (ferritin light chain – Chromosome 19), *TMSB10* (thymosin beta 10 – Chromosome 2), *TMSB4X* (thymosin beta 4 X-linked – Chromosome X), *GAPDH* (glyceraldehyde-3-phosphate dehydrogenase – Chromosome 12), and *ALDH1A1* (Aldehyde dehydrogenase – Chromosome 12), respectively. We classified these genes into low, medium and high expression levels based on the published results from whole transcriptome direct cDNA sequencing in the well-characterized A549 cell line^9^ (**Fig. 1A, Table S1**). We designed all CRISPR RNAs (crRNAs) to bind to the first exon at 5’ ends or the last exon at 3’ ends and generate a single end cut for ligating sequencing adapters (**Fig. 1B, Methods, Table S2**). crRNAs were pooled before Cas9 nuclease cutting and adaptor ligation for sequencing.

**Figure 1:**
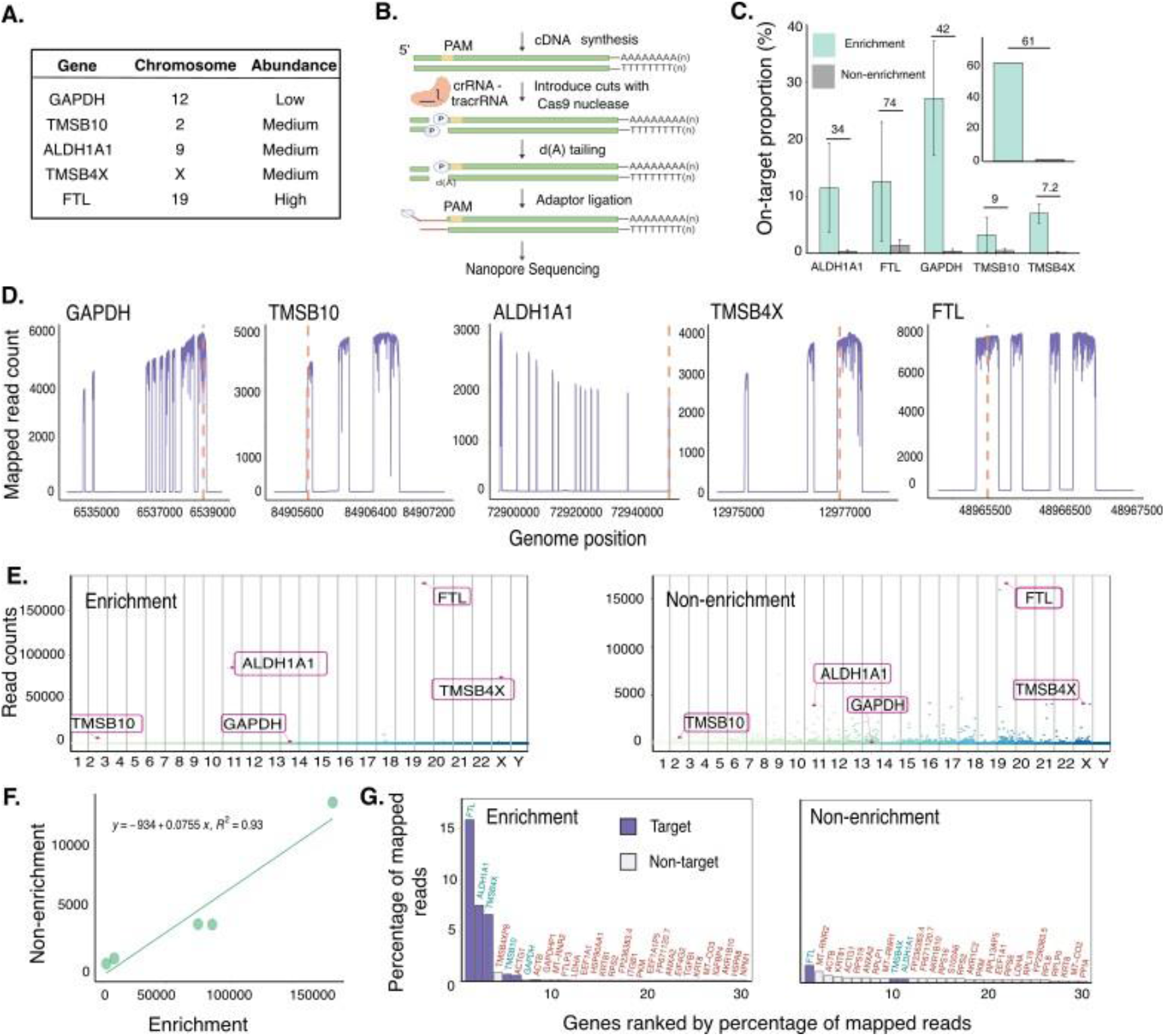
Enrichment of human transcript in A549 cell line. **(A)** Five target transcripts with wide range of expression. **(B)** Cas9-targeted sequencing: cDNA is synthesized by using a poly(T) primer. gRNA binds to the target site allowing Cas9 nuclease cutting. Adapters are ligated to phosphate groups on both strands of the cDNA. **(C)** The proportion of mapped reads in each transcript in enriched and non-enriched cDNAsequencing experiment. An inset shows the total target proportion. Cas9-targeted sequencing significantly increases the proportion of target transcripts among the transcriptome (*Two-tailed t-test, p=0.0002*) (**Table S3**). **(D)** Alignment coverage of each transcript. All transcripts have even coverage in all known exons. Dashed red lines indicate gRNA binding sites. **(E)** Read counts at the gene level of all mapped reads in enrichment and non-enrichment sequencing. Each dot represents the read count of specific transcript. Columns are the human chromosomes. Cas9-targeted sequencing reduces sequencing noise compared with whole transcriptome sequencing and promotes the identification of target transcripts. (**F)** Highly correlated expression between enrichment and non-enrichment sequencing at gene level. **(G)** 30 highly abundant transcripts in enriched and unenriched samples. Red color indicates off-targets, while blue color represents on-target transcripts.

We compared our enrichment data to the published direct cDNA sequencing of the same cell lines^9^. Our approach achieved significantly higher coverage across all on-target transcripts by using approximately 1μg of poly(T)_28_ synthesized cDNA samples and running on MinION flow cells (both R9.4.1 and R10.4.1 versions) (*Two-tailed t-test, p=0.0002*) (**Fig 1C, Methods, Table S3**). On average, target mapping coverage ranged from 330x to 2588x (**Table S4**). Gene exhibiting the highest fold change of coverage was *FTL* while *TMSB10 s*howed the lowest enrichment with a 7.2- fold change (**Fig. 1C**). Reference lengthsof the targets range from 467 bp to 2096 bp,slightly longer than the majority read length generated by our method, which is due to the Cas9 cutting. Despite this, our method generated even depth for all exons of the targets (**Fig. 1D**).

We then assigned all mapped reads to genomic features and evaluated the expression of each target at the gene level. In total, more than 50% of feature-assigned reads were on-target, compared to less than 1% in unenriched sequencing (**Table S4**). In each target chromosome, Cas9 enrichment amplified the target while reducing noise caused by other high expression genes, which were abundantly found in unenriched sequencing (**Fig. 1E**). The expression of the targets was highly correlated with the estimated abundances by whole transcriptome cDNA sequencing (*Pearson’s correlation of 0.9493*, **Fig. 1F**). This result indicates that Cas9-targeted sequencing can effectively preserve the relative proportions of the target transcripts.

Next, we investigated off-targets in our experiment by extracting the 30 most abundant transcripts. We considered two possibilities: if most of the off-targets were found in both enrichment and non-enrichment methods and expressed at high levels, this would indicate the random ligation of the adapters; on the other hand, we would find the enrichment of the off-targets which are only available in Cas9-enrichment data. We found that 13 out of 25 (52%) of the off-targets were also present among the most abundant transcripts in non-enrichment sequencing data (**Fig. 1G, Table S5**). This suggests that off-targets are primarily due to random ligation, which is similar to that observed in Cas9-targeted sequencing to enrich chromosomal DNA25.

### Cas9-targeted sequencing amplifies target signals for isoform identification

Since we obtained substantial high depth for target transcripts in the previous experiment, here, we applied the Cas9-targeted approach to focus on low abundance mRNA with well-known isoforms. KPNA2 is a member of the nuclear transporter family and is involved in the delivery of numerous cargo proteins from the cytoplasm to the nucleus, which has been showed to contribute to cancer development^26,27^. Several predicted isoforms have been reported in which *ENST00000300459 (KPNA2-201)* and *ENST00000537025 (KPNA2-202)* were highly expressed and showed differential transcript usage during viral infection in specific cell types^9^. The two variants are distinct in their lengths, particularly in the first exon (**Fig. 2A**). Thus, we designed a gRNA binding to the second exon which generates two fragments, a long fragment including a portion of exon 2 and exon 3-11; and a short fragment including exon 1 and a part of exon 2 (**Fig. 2D**).

**Figure 2:**
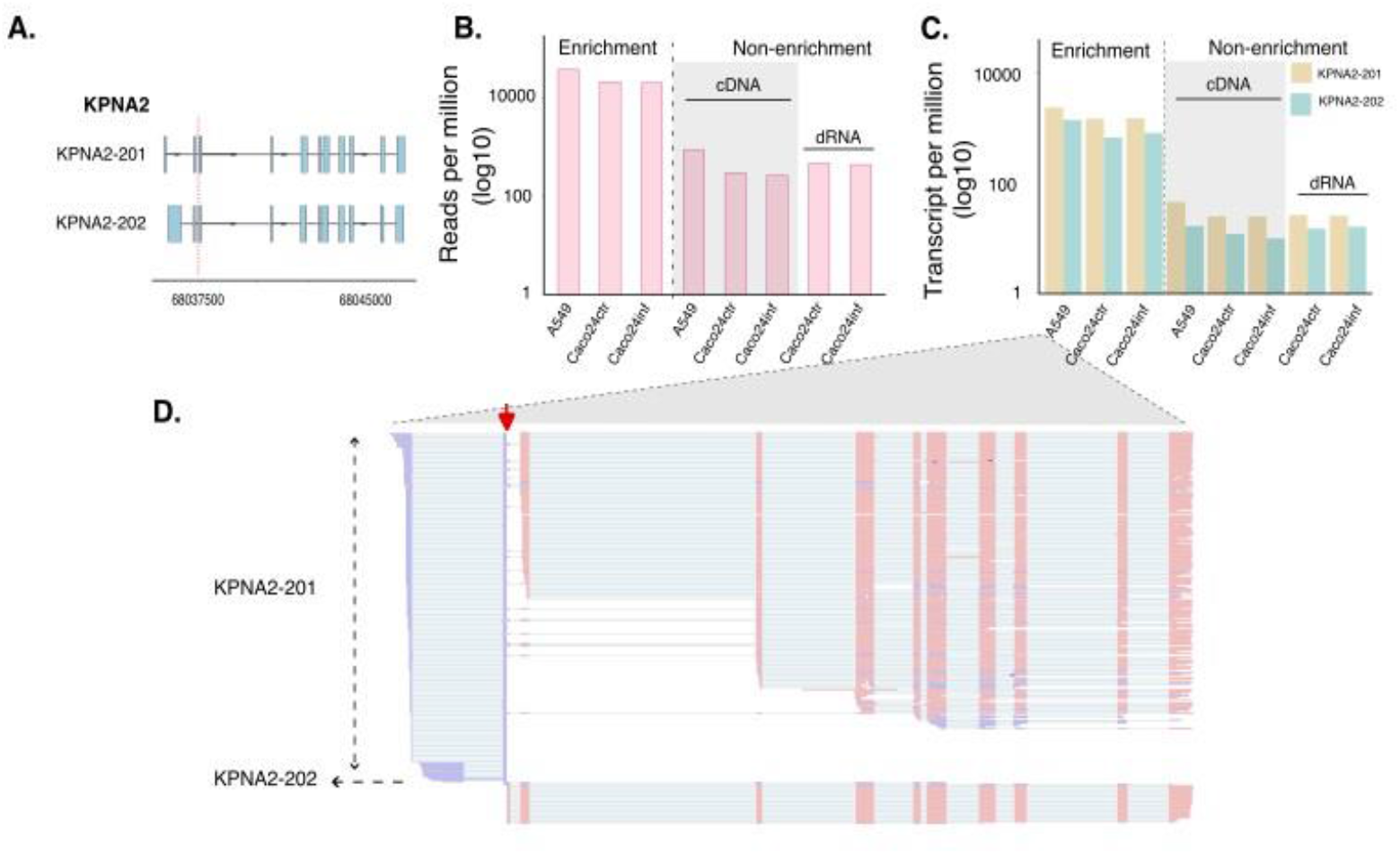
Identification of *KPNA2* isoforms using targeted sequencing. **(A)** Two highly expressed transcripts in human cell lines. **(B)** Gene level counts for KPNA2 in enrichment and non- enrichment samples. We obtained higher counts for enrichment samples compared to whole transcriptome cDNAand dRNAsamples. **(C)** Isoform expression in infected and non-infected cells. There were no differences in isoform abundance between infected and non-infected cells. **(D)** IGV (Integrative genomics viewer) screenshot displaying mapping coverage of KPNA2. Reads were colored by read strand, blue for negative strand, pink for positive strand. Red arrow indicates the Cas9 nuclease cutting site.

We first re-analyzed whole transcriptome data generated by both direct cDNA sequencing and dRNA-seq from previous study, which utilized the same A549 and Caco-2 cell lines^9,28^ (**Method**). In these cell lines, the *KPNA2* gene expressed at comparably low levels. Particularly, at gene level, only 0.09% of reads mapped to this transcript in the A549 sample. Similarly, Caco-2 control and Caco-2 infected with SARS-CoV-2 samples expressed the *KPNA2* gene at 0.03%. By using the Cas9-targeted system, we obtained a significant increase in *KPNA2* coverage (*Two-tailed t-test, p=0.0008*) (**Fig. 2B**), an average of 62-fold higher coverage of *KPNA2* transcript compared to unenriched samples.

To identify the proportion of the isoforms in these samples, we quantified the isoform expression by using *Salmon* (**Methods**). In alignment with previous study^9^, we found that *KPNA2-201* is dominantly detected in both Caco-2 and A549 cell lines, and the proportion of these isoforms were preserved across enrichment and non-enrichment methods (**Fig. 2C, Table S6**). Interestingly, we were able to detect the reads that cover exon 1 of *KPNA2-202*, which is rarely seen in cDNA sequencing and full-length dRNA-seq (**Fig. 2D, Fig. S1**).

### Enrichment of SARS-COV-2 RNA improves the identification of diverse viral sub-genomes in infected cells

Viral infections, such as those with SARS-CoV-2, cause a broad spectrum of disease states with widely variable severity and symptoms. Deep genomic sequencing is critical for early treatment and understanding the viral distribution and circulation *in vivo*. However, it is difficult to obtain high depth sequencing due to the high ratio of host versus pathogen RNA and the potential for RNA degradation, especially in lowly infected tissue. Thus, we applied Cas9 enrichment to target both human and viral transcripts. The SARS-CoV-2 genome is a singular, non-segmented large positive sense-stranded RNA with a length of about 30 kb. From the genome, a set of plus-strand sub- genomic RNAs (sgRNAs) are generated with a common leader sequence of 70 nucleotides located at 5’-end and a section at the 3’ end of the genome^29^. These sgRNAs encode the following viral proteins: structural proteins - spike (S), envelope (E), membrane (M) and nucleocapsid protein (N) and several accessory proteins - 3a, 6, 7a, 7b, 8, and 10. To determine the efficiency of Cas9-targeted sequencing in capturing a small fraction of SARS-CoV-2 transcriptome in the samples with low viral load, we designed a crRNA that targets the 5’- end leader sequence to capture all possible viral structural transcripts and accessory transcripts (**Fig. 3A, Methods**). We then compared our enrichment data with data generated from whole transcriptome dRNA-seq and cDNA sequencing of the same infected samples.

**Figure 3:**
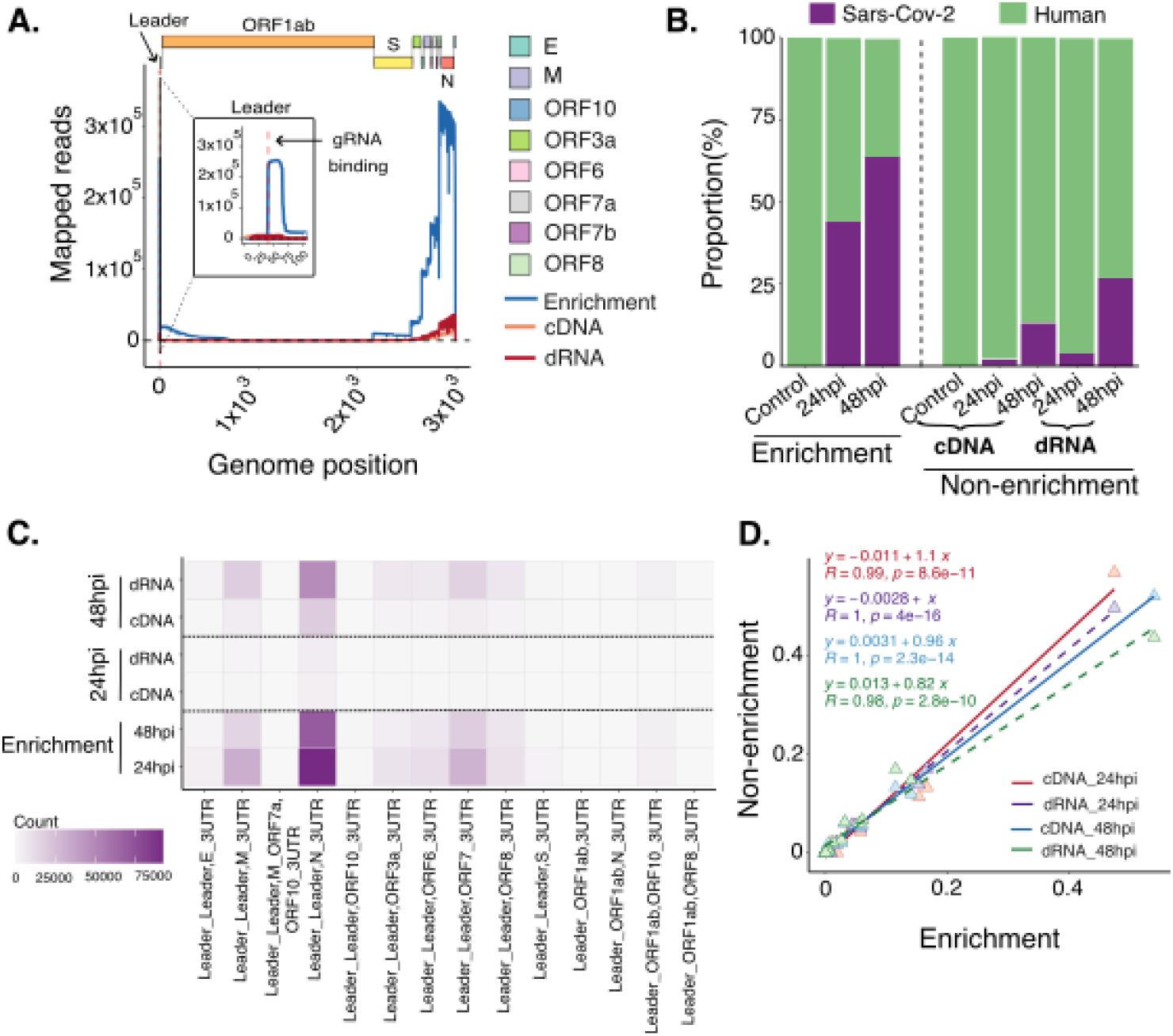
Enrichment of SARS-CoV-2 transcripts in human infected cells. **(A)** SARS-CoV-2 genome alignment. Mapped reads covered all ORFs with substantial reads mapping at 5’ leader sequence and all sub-genome ORFs: enriched (blue), unenriched: cDNA (orange) and dRNA-seq (red) **(B)** Proportion of human reads and viral reads. While unenriched samples maintained high human proportions, the enrichment method increased the proportion of viral reads. **(C)** The abundance of different classes of sgRNA in enriched and unenriched samples of infected cells at different time points. Cas9 enrichment method recovered all transcripts with higher coverage compared to whole transcriptome cDNA and dRNA sequencing. **(D)** Highly correlated abundance of sub-genome transcripts between Cas9 enrichment method and bulk sequencings (*Pearson’s correlation*).

We first mapped all enriched reads to the SARSCoV-2 and human genome to identify the fraction of viral reads in the samples (**Methods**). We were able to cover the whole genome of SARS-CoV-2 (**Fig. 3A**). Particularly, in Caco-2 infected cells harvested 48-hour post-infection (hpi), the median read length was 4972 nt, and 63 reads were mapped to the genome with lengths longer than 25 kb. We observed high coverage for the region of the leader sequence and at the 3’ end of the viral genome (**Fig. 3A**). Moreover, we noted a difference in the length of the leader sequence between enriched and unenriched samples due to the cutting of Cas9 nuclease. There was uniform coverage across the body of each annotated open reading frame (ORF), with a sharp drop at the end of the ORF. This observation is consistent with the previous studies and supports the accepted model of SARS-CoV-2 discontinuous transcription, where most of the viral sgRNAs are the result of the fusion of an ORF with the 5’ leader sequence^29^.

We then identified the proportion of viral reads compared with human reads. We found a significant increase in viral reads compared to non-enrichment methods (*Two-tailed t-test, p=0.0156*). Particularly, more than 40% and 60% of generated reads were mapped to the SARS-CoV-2 reference genome while there were only 2% and 15% of reads across all non-enrichment samples with 24 hpi and 48 hpi, respectively (**Fig. 3B**). We did not detect any viral reads in control sample.

Next, we assigned all mapped reads to specific viral sgRNAs to examine the transcript expression (**Methods**). We obtained significantly higher read counts for each sgRNA compared to non-enrichment samples and covered all sgRNAs that were found in both direct cDNA sequencing and dRNA-seq (**Fig. 3C**). Our enrichment data was highly correlated with whole-transcriptomic sequencing data (**Fig. 3D**) and exhibited a similar expression pattern of structural and accessory proteins with nucleocapsid N as the most highly expressed (**Fig. 3C**).

## Discussion

Full-length RNA sequencing facilitates the identification of transcript isoforms. However, the limitation is that low abundance transcripts are often missed due to the lack of depth. Targeted transcript long-read sequencing leverages the ability to sequence transcript molecules end-to-end, while circumventing the weakness of limited sequencing yield and low transcript coverage. Although, there are existing solutions for targeted long-read RNA sequencing, these are either associated with high cost, or complicated to set up and implement. Here, we employed targeted CRISPR-Cas9 sequencing to enrich the preselected transcripts among transcriptomes. We tested the method in several cell lines with wide range of transcript abundance. Our comprehensive benchmark analysis indicated consistently high on-target rate and fold enrichment across all samples. We showed that Cas9-targeted sequencing substantially improved the sensitivity of detecting low- abundance transcripts as well as the identification of these transcript isoforms. At the same time, the estimated abundances of target transcripts highly correlated with the whole transcriptome cDNA sequencing and dRNA-seq. We also showed that Cas9-targeted sequencing was able to enrich transcripts across species by testing the method on SARS-CoV-2 infected cell lines in different time points. Cas9-targeted sequencing provided clear evidence of the expression of viral accessory transcripts which were missed out in bulk cDNA sequencing. Overall, these results indicate that Cas9-targeted sequencing provides a robust tool for the quantification for specific transcripts as well as transcript isoform confirmation and possibly new isoform discovery.

Cas9-targeted sequencing facilitates many applications of targeted long-read RNA sequencing. Here, we illustrated the application of this Cas9 method to enrich a small scale of gene panels and only focused on human and viral transcripts. However, Cas-9 targeted sequencing can be applied to any gene panel of interest for focused discovery and quantification of transcript isoforms. We anticipate that the strategy for applying Cas9-targeted sequencing will have a broad impact, facilitate the significant cost reduction as well as less ease of implementation, and therefore will pave the way for wide adaptation in diverse biomedical and clinical settings.

## Methods

### Cell culture and viral infection

For this study, all cell lines were maintained at 37°C with 5% CO_2_. The culture of A549 (human lung carcinoma epithelial) was performed in F12 media (Gibco) supplemented with 10% FCS, GlutaMAX, and Sodium Pyruvate. Cell culture of Caco-2 (human intestinal epithelial cells, ATCC Cat#HTB-37; RRID: CVCL_0025) has been described in previous studies^9,28^. Briefly, Caco-2 cells were cultured in Dulbecco’s Modified Eagle Medium (DMEM) (Media Preparation Unit, Peter Doherty Institute) supplemented with 1X non-essential amino acids (Sigma-Aldrich), 20 mM HEPES, 2 mM L-glutamine, 1X GlutaMAX 2 μg/mL Fungizone solution, 26.6 μg/mL gentamicin, 100 IU/mL penicillin, 100 μg/mL streptomycin, and 20% FBS. For viral infection, the cell line was seeded in 4 × 6-well tissue-culture plates and infected with SARS-CoV-2 (Australia/VIC01/2020) virus at a multiplicity of infection (MOI) of 0.1. The infected cells were harvested at 2, 24 and 48 hour post-infection (hpi)

### RNA extraction, cDNA synthesis and purification

For this study, cultured cells of A549 were harvested by centrifuge and extracted RNA following manufacturer’s instructions for the RNeasy Mini Kit (QIAGEN). All extracted RNA was then treated with DNase from the Turbo DNA-free kit (Invitrogen) to reduce DNA contamination and purified by RNAclean XP magnetic beads (Beckman Coulter). Maxima H minus Double-Stranded cDNA synthesis (Thermoscientific – K2561) was used to synthesize cDNA for Cas9-targeted sequencing. The concentration and fragmentation of extracted RNA and synthesized cDNA were measured by Nanodrop 2000C (Thermo Fisher Scientific), 4200 TapeStation System (Agilent Technology), and Qubit 4 Flourometer (Invitrogen). RNA samples of control and infected cells harvested at 24 and 48 hpi from Caco-2 were obtained from our previous study with similar sample preparation protocol^9^.

### Cas9-targeted sequencing

*Design of gRNAs*: All crRNAs were designed using CHOPCHOP (version 3) web tool to select target sites specific for CRISPR/Cas9 nanopore enrichment (https://chopchop.cbu.uib.no/about). Human genome (Homo sapiens – hg38/GRCh38) and SARS-CoV-2 genome (Australia/VIC01/2020) served as the references. The sequences of crRNAs are provided in **Supplementary TableS2**.

### Cas9 cleaving and library preparation

The preparation for Cas9 nuclease cutting and library preparation for sequencing followed the recommended protocol from Oxford Nanopore Technologies with some modifications. In brief, all crRNAs were pooled and combined with tracrRNA (trans-activating crRNA) in Duplex Buffer to generate gRNAs. HiFi Cas9 Nuclease V3 (IDT) was mixed with gRNAs in 1x CutSmart buffer (NEB) to generate Cas9 ribonucleoprotein complex (RNP). Approximately 1 μg of pooling synthesized cDNAwas dephosphorylated by Quick calf intestinal alkaline phosphatase (CIP). Dephosphorylated cDNA was cut by RNP and dA-tailing by NEB Tag polymerase followed by library preparation. Ligation sequencing Kits including SQK- LSK109 and SQK-LSK114 were utilized to prepare sequencing library for R9.4.1 and R10.4.1 flow cells, respectively. Samples were then run on a MinION or GridION sequencer. A detailed description of runs (flow cell, gRNA, sequencer) is provided in **Supplementary Table S1**.

### Sequencing analysis

*Basecalling and read alignment of nanopore sequencing data:* For both R9.4.1 and R10.4.1 chemistry samples, *Dorado* (v0.7.0) was used to perform the basecalling. Model *dna_r9.4.1_e8_sup@v3.6* was applied to R9.4.1 samples and model *dna_r10.4.1_e8.2_400bps_sup@v5.0.0* was applied to R10.4.1 samples. Aparameter “--min-qscore 7” was set to filter out low quality reads. *Samtools* (v1.16.1) was used to convert the output ubam to fastq. Reads were mapped to human reference genome (Homo_sapiens GRCh38 release 100) and SARS-CoV-2 genome (SARS-CoV-2/human/AUS/VIC01/2020) using *Minimap2* (v2.24) with parameter “-y --eqx -uf -ax splice -k 12” for human and “-aL –x map-ont” for SARS-Cov-2. ON-target reads were defined as those that aligned within the start position and end position of a gene. Average coverage and read length analyses were obtained by *Samtools*.

For whole transcriptome direct cDNA and dRNA-seq, the data for A549, Caco-2 mock control, and Caco-2 infected with SARS-CoV-2 harvested after 24 and 48 hpi was obtained from previous study^9^. *Guppy* (v3.5.2) was used to perform the basecalling with configuration file as *dna_r9.4.1_450bps_hac.cfg*. Similarly, reads alignment, coverage and read length analysis were performed using the same methods as previous analysis.

### Transcript analysis

For human, gene-level read counts were generated by *FeatureCounts*^*30*^ (v2.0.6). For SARS-CoV-2, our in-house tool, *npTranscript* (v1.0 - *https://github.com/lachlancoin/npTranscript*), was utilized to assign read to genome and sub- genome transcripts. *Salmon*^*31*^ (v1.10.1) was used for isoform quantification with parameter “-- noErrorModel -l U” with transcriptome reference (GENCODE release 35). For each sample, we calculated an on-target rate by dividing the number of reads mapped to target by the total number of reads mapped to all features of the reference genome. Fold enrichment on a given sample was calculated by dividing the on-target rate for Cas9-targeted sequencing by the on-target rate for the non-enrichment sample.

### Statistics and illustration

All statistics and plots were performed by using *GraphPad Prism* (v10.4.1) for MacOS and *Rstudio* (v2024.04.2).

## Data availability

Sequencing data for this study can be found in NCBI with accession number PRJNA1244050 Data for direct cDNA sequencing and dRNA-seq obtained from previous study can be found in online repositories: 1. https://www.ncbi.nlm.nih.gov/, PRJNA675370. 2. https://figshare.com/, 10.6084/m9.figshare.17139995. 3. https://figshare.com/, 10.6084/m9.figshare.16841794. 4. https://figshare.com/, 10.6084/m9.figshare.17140007.

## Notes

### Competing Interest Statement

LC has received travel funding and research funding from Oxford Nanopore Technologies unrelated to this manuscript.

### Funding

- National Health and Medical Research Council: GNT1195743

